# Impaired cerebellar plasticity hypersensitizes sensory reflexes in *SCN2A*-associated ASD

**DOI:** 10.1101/2023.06.05.543814

**Authors:** Chenyu Wang, Kimberly D. Derderian, Elizabeth Hamada, Xujia Zhou, Andrew D. Nelson, Henry Kyoung, Nadav Ahituv, Guy Bouvier, Kevin J Bender

**Affiliations:** Neuroscience Graduate Program, University of California, San Francisco, San Francisco, CA, USA; Department of Neurology, University of California, San Francisco, San Francisco, CA, USA; Weill Institute for Neurosciences, University of California, San Francisco, San Francisco, CA, USA; Department of Bioengineering and Therapeutic Sciences, University of California, San Francisco, San Francisco, CA, USA; Institute for Human Genetics, University of California, San Francisco, San Francisco, CA, USA; Department of Physiology, University of California, San Francisco, San Francisco, CA, USA

## Abstract

Children diagnosed with autism spectrum disorder (ASD) commonly present with sensory hypersensitivity, or abnormally strong reactions to sensory stimuli. Such hypersensitivity can be overwhelming, causing high levels of distress that contribute markedly to the negative aspects of the disorder. Here, we identify the mechanisms that underlie hypersensitivity in a sensorimotor reflex found to be altered in humans and in mice with loss-of-function in the ASD risk-factor gene *SCN2A*. The cerebellum-dependent vestibulo-ocular reflex (VOR), which helps maintain one’s gaze during movement, was hypersensitized due to deficits in cerebellar synaptic plasticity. Heterozygous loss of *SCN2A*-encoded Na_V_1.2 sodium channels in granule cells impaired high-frequency transmission to Purkinje cells and long-term potentiation, a form of synaptic plasticity important for modulating VOR gain. VOR plasticity could be rescued in adolescent mice via a CRISPR-activator approach that increases *Scn2a* expression, highlighting how evaluation of simple reflexes can be used as quantitative readout of therapeutic interventions.

## INTRODUCTION

Altered sensitivity to sensory input is a hallmark of autism spectrum disorder (ASD); over 90% of children with ASD experience heightened sensitivity to sensory stimuli, spanning sensory modalities (*1*–*4*). Although the neuronal mechanisms that contribute to this hypersensitivity have been examined extensively in forebrain circuits (*5*–*8*), emerging evidence suggests that changes in sensory sensitivity can occur even at much earlier stages of sensory processing, including those that support sensory reflexes (*9*–*12*). Thus, changes in reflexive behavior can provide a window into cellular and circuit dysfunction that underlie altered sensory experience in ASD.

The vestibulo-ocular reflex (VOR) is one such reflex that is affected in ASD. This reflex transforms vestibular sensory information generated by head movements in one direction into eye movements in the opposite direction, thereby helping stabilize images on the retina. In neurotypical subjects, this reflex is engaged well by fast head movements, but not by slower head movements. By contrast, VOR assessed in children with ASD appears to be sensitive to both fast and slow movements, reflecting a hypersensitivity not seen in neurotypical children (*10, 12*). The mechanisms for this heightened sensitivity are unknown.

Here, we show that VOR gain is hypersensitized in children and mouse models of *SCN2A* loss-of-function (LoF), a condition that carries the highest risk for ASD of any gene identified via clinical exome sequencing (*13, 14*). *SCN2A* encodes the neuronal sodium channel Na_V_1.2 (*15*). This channel supports action potential (AP) initiation and propagation in multiple cell classes (*16*–*19*), including cerebellar granule cells whose activity is key for modulating VOR gain (*20*). *SCN2A* LoF impaired plasticity between granule cells and Purkinje cells, in turn hypersensitizing VOR by preventing synaptic plasticity that typically readjusts VOR amplitude (*21*–*24*). Remarkably, VOR plasticity could be restored in adolescent mice by upregulating *Scn2a* expression levels via a CRISPR-activator (CRISPRa) based approach. Overall, these data demonstrate how innate reflexes provide a window into cerebellar dysfunction in ASD that is both well-conserved across species and sensitive to therapeutic intervention.

## RESULTS

### *SCN2A* loss-of-function hypersensitizes VOR gain in humans and mice

Children with *SCN2A* LoF variants typically have limited to no verbal repertoire and have difficulty following instructions, but often are very comfortable with caregivers (*15*). With these behavioral aspects in mind, we developed a lightweight, helmet-mounted infrared eye-tracking system paired with an inertial measurement unit (IMU) to assess both eye and head movement, respectively (Fig. 1A, S1). Children aged 3-10 years were seated either alone or on a caregiver’s lap, and VOR gain was assessed with ±5° sinusoidal head oscillations in the dark (0.4 Hz, peak angular velocity: 12.5°/s). Consistent with previous work, VOR gain was well below unity in neurotypical children (*10*). By contrast, VOR gain was near unity in children with *SCN2A* LoF variants (Fig. 1B, E).

**Figure 1:**
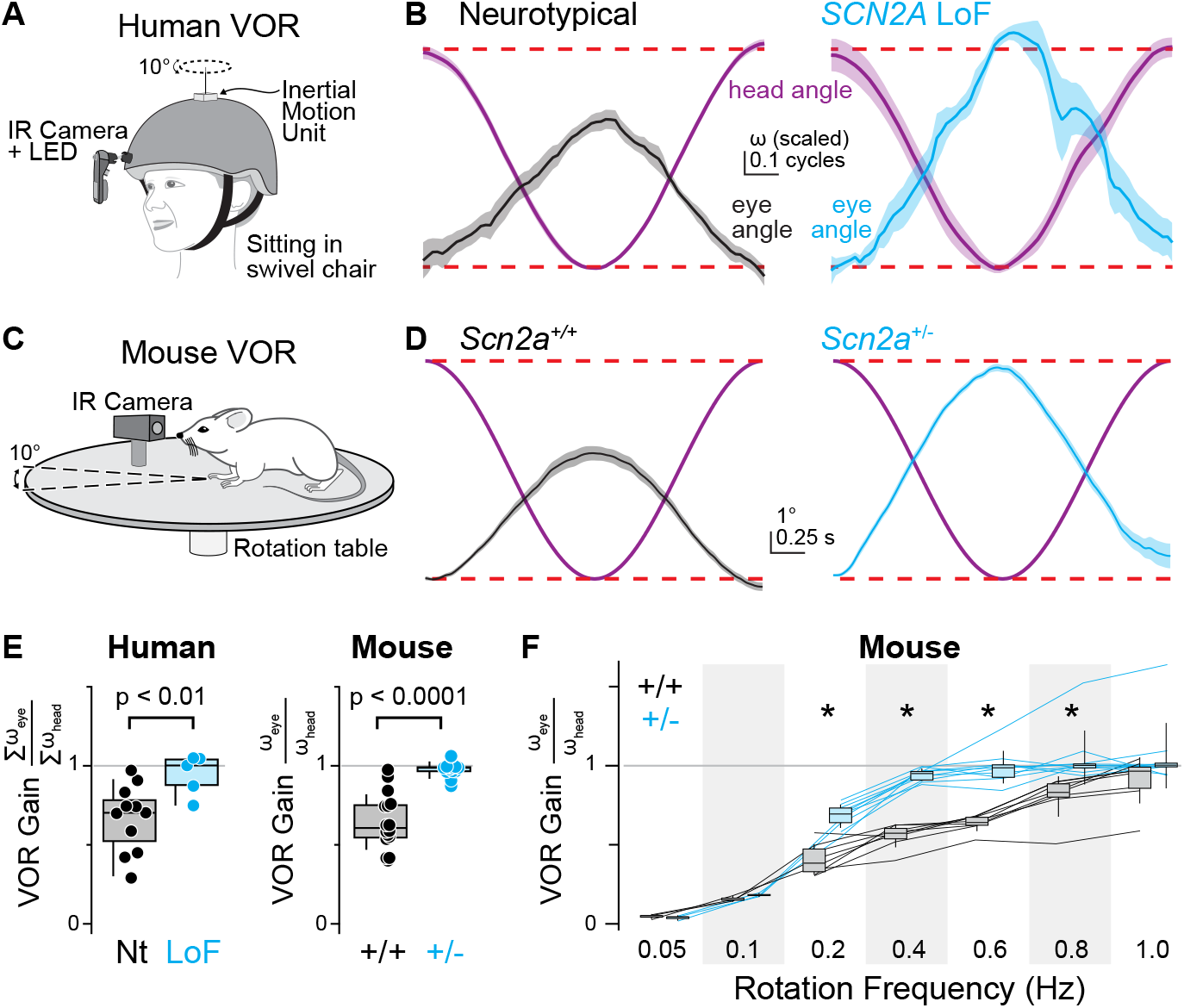
VOR gain is elevated in *Scn2a* haploinsufficiency conditions. **A:** Schematic of purpose-built eye tracking apparatus for VOR assessment in children. Head movements are measured from a device located at the center of the head while the right eye is imaged under infrared illumination (940 nm LED). Participants are seated in a swivel chair and oscillated ±5° at ∼0.4 Hz to assess VOR gain. **B:** Head angle (purple) and contraversive eye angle for neurotypical (Nt, black) and *SCN2A* LoF conditions (cyan). Lines represent the average of a single cycle from all participants; shaded area is SEM. Dashed red lines indicate ±5° range. **C:** VOR was assessed in mice head-fixed at the center of a rotating table, imaged under IR illumination. **D:** VOR at 0.4 Hz rotation frequency, displayed as in panel B, for *Scn2a*^*+/+*^ (left, black) and *Scn2a*^*+/-*^ mice (right, cyan). **E:** Baseline VOR gain in human and mouse. Circles are individuals; box plots are medians, quartiles and 90% tails. n: 11 Nt, 5 LoF humans; 13 +/+, 12 +/-mice. Mann Whitney test p-values shown. **F:** VOR gain across rotation frequencies in mouse. Lines connect repeated tests in single mice. Asterisks: p < 0.0001, Friedman test on overall distribution; Mann Whitney test on individual frequencies, Holm Šídák correction.

Protein truncation and resultant nonsense-mediated decay accounts for the vast majority of *SCN2A* LoF cases (*13, 15*). As such, mice heterozygous for *Scn2a* are an ideal model system for studying LoF-related physiology. We therefore assessed VOR in *Scn2a*^*+/-*^ mice head-fixed on a rotational platform using identical test conditions as above (±5° rotation at 0.4 Hz in the dark). Like children with *SCN2A* LoF, *Scn2a*^*+/-*^ mice showed a VOR gain near unity, significantly higher than wild type (WT) littermates. This elevated gain persisted over a wide range of rotation frequencies (Fig. 1F). This suggests that aberrantly high VOR gain may serve as a readout for *SCN2A* LoF both in mouse and human.

### VOR hypersensitivity is associated with an inability to reduce VOR gain

VOR gain is plastic and is adaptively increased or decreased throughout life to maintain image stability on the retina. The high gain in humans and mice suggests that VOR plasticity has been compromised, and that *SCN2A* LoF impairs one’s ability to reduce VOR gain. To test this, we implemented a standard protocol that decreases VOR gain, termed “VOR gain-down”. Here, instead of rotating the animal in the dark, the animal was presented with a visual stimulus of high-contrast vertical bars, displayed on surrounding computer monitors, that moved in concert with the animal (Fig. 2A). During induction, both WT and *Scn2a*^*+/-*^ mice tracked the visual stimulus, completely suppressing VOR despite the animal experiencing head movement (Fig. S2A, C). Typically, this strong VOR cancellation during the induction protocol results in long-lasting reductions in VOR gain, assessed again in the dark right after induction. This persistent reduction in gain was indeed observed in WT mice. By contrast, VOR gain, when assessed in the dark after induction, immediately returned to unity in *Scn2a*^*+/-*^ mice (Fig. 2B-D). Thus, in *Scn2a*^*+/-*^ mice, VOR gain is elevated and is not reduced by a gain-down protocol.

**Figure 2:**
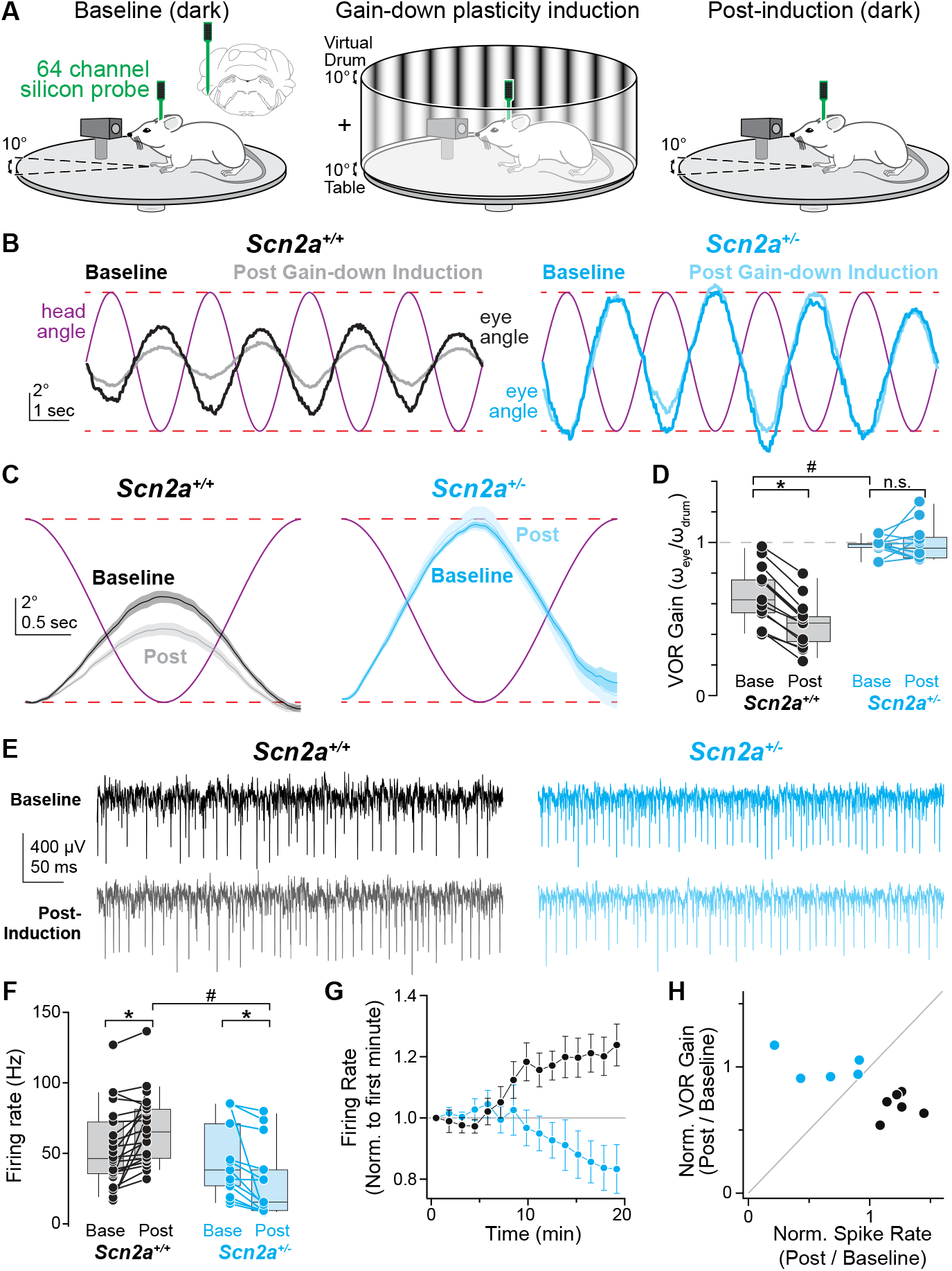
Impaired VOR gain-down behavioral plasticity in *Scn2ar*^*+/-*^ mice. **A:** Experimental design for VOR gain-down induction and simultaneous *in vivo* silicon probe recording in the cerebellar floccular complex. Inset, recording location in left hemisphere. Left: baseline VOR recording in the dark. Middle: VOR gain-down induction with virtual drum as visual stimulus. During gain-down induction, visual stimulus was moved in phase with table. Right: post-induction VOR recording in the dark. **B:** Head and eye angles in *Scn2a*^*+/+*^ (black) and *Scn2a*^*+/-*^ (cyan) during baseline (dark colors) and post-induction (light colors) with table rotation (purple). Dashed red lines indicate ±5° range. **C:** Head angle (purple) and contraversive eye angle before (darker shade) and after (lighter shade) gain-down induction. Data presented as in Fig. 1D. **D:** VOR gain before and after gain-down induction. Data color coded as in B. Bars connect data from individual mice. * p < 0.001, Wilcoxon signed rank test, # p < 0.0001, Mann Whitney test. **E:** Putative Purkinje cell unit recordings before and after gain-down induction. Traces are aligned from table rotation onset. **F:** Average Purkinje cell simple spike firing frequency during sinusoidal head rotation, before and after gain-down induction, displayed as in D. *Scn2a*^*+/+*^ (black, closed circles, n = 22 units, 6 mice), *Scn2a*^*+/-*^ mice (cyan, n = 13 units, 5 mice). * p < 0.05, all conditions, Mixed-effects modeling, # p < 0.001, Wilcoxon signed rank test. **G:** Normalized Purkinje cell simple spike firing frequency during induction protocol, normalized to firing rate in first minute per unit. Circles and bars are mean ± SEM, binned per minute. **H:** Normalized change in VOR gain vs. normalized change in simple spike rate (normalized per unit and averaged across units per animal). Circles are color-coded as in panel D and represent individual animals.

In the cerebellum, the floccular complex is critical for modulating VOR gain (*25, 26*). VOR plasticity has been proposed to be mediated in part by the modulation of Purkinje cell activity (*22, 27, 28*). To test whether the firing rate of Purkinje cells changes following VOR gain-down induction in *Scn2a*^*+/-*^ mice, we used silicon probes to monitor extracellular activity from the left floccular complex before, during, and after gain-down induction, contralateral to the imaged eye (Fig. 2, S3). Putative Purkinje cell units were modulated during head rotation in both WT and *Scn2a*^*+/-*^ mice, on average preferring clockwise head rotations over counterclockwise rotations (Fig. S4). In addition, Purkinje cell firing rates were increased following gain-down induction in WT mice (Fig. 2E-F). Firing rate changes occurred within 10 minutes of induction onset (Fig. 2G) and were consistent across all animals studied (Fig. 2H). But in *Scn2a*^*+/-*^ mice, Purkinje cell activity did not increase following gain-down induction. Instead, firing rate decreased modestly, indicating that normal plasticity mechanisms were impaired in *Scn2a*^*+/-*^ mice (Fig. 2E-H).

### *Scn2a* heterozygosity alters granule-to-Purkinje cell synaptic transmission and plasticity

Within the cerebellum, Na_V_1.2 expression appears limited to granule cells (Fig. 3A-C) (*29*–*31*), with highest density in their parallel fiber axons that make excitatory synapses onto Purkinje cells. Gain-down plasticity is correlated with long-term potentiation (LTP) of parallel fiber-to-Purkinje cell synapses (*32*–*34*). Thus, deficits in granule cell function in *Scn2a*^*+/-*^ mice may affect synaptic plasticity that mediates VOR plasticity. To test this, we made acute slices of cerebellum and stimulated parallel fibers with a high-frequency burst-based LTP protocol and monitored EPSC amplitude and paired-pulse ratio (PPR) in Purkinje cells. This protocol evoked robust potentiation of EPSCs in WT mice (Fig. 3D), and was correlated with a reduction in PPR, suggesting that this induction protocol acts, at least in part, to increase release probability (*35, 36*). By contrast, no LTP was observed in *Scn2a*^*+/-*^ mice, and PPR was unchanged (Fig. 3D). Thus, a lack of VOR gain-down plasticity may be due to difficulty in potentiating synapses between granule and Purkinje cells in *Scn2a*^*+/-*^ mice.

**Figure 3:**
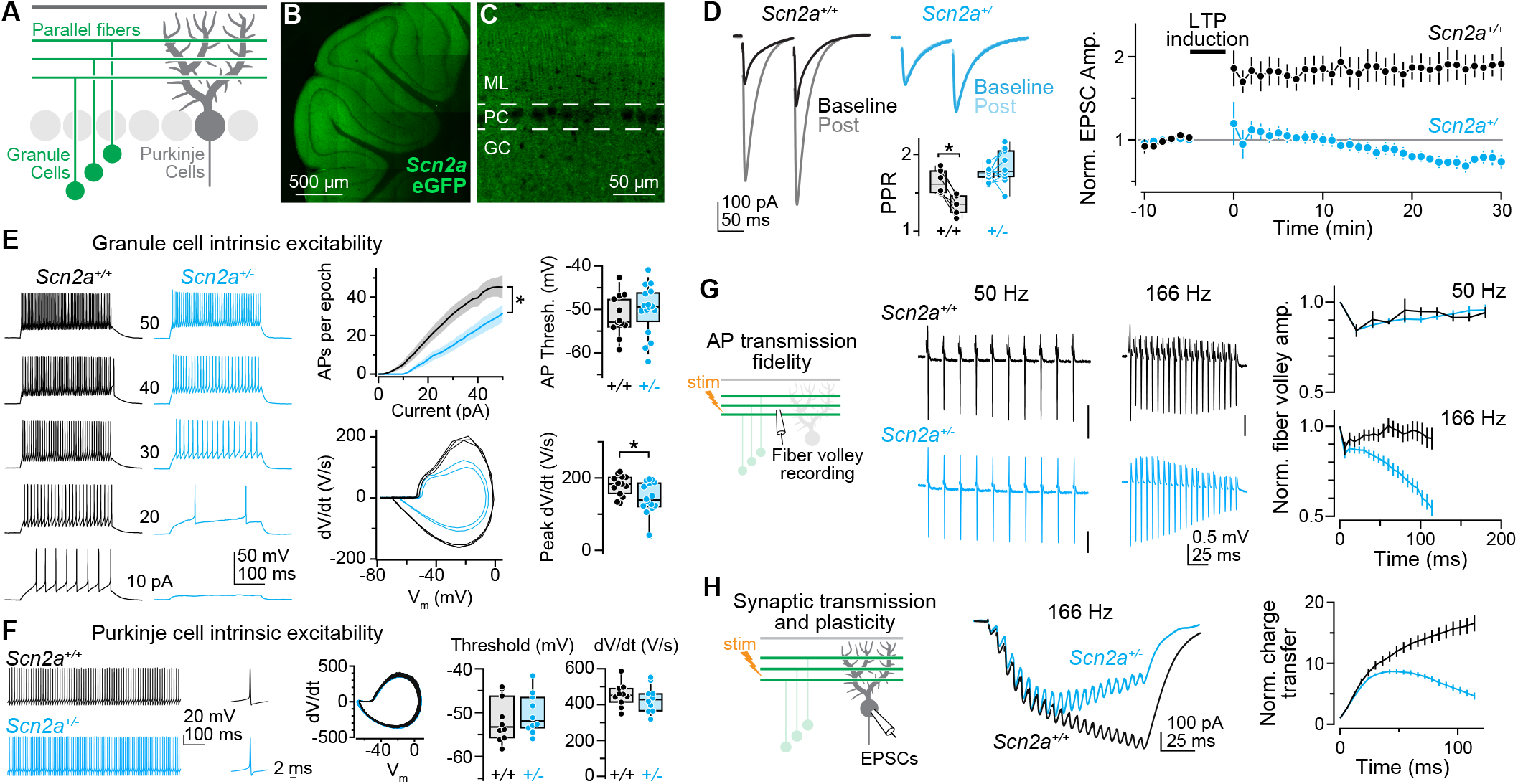
Impaired cerebellar granule cell excitability and plasticity in *Scn2ar*^*+/-*^ conditions. **A:** Schematic of excitatory circuit between granule cells and Purkinje cells. Subsequent panels highlight different circuit components under investigation. **B:** Coronal section of the mouse cerebellum in animal with eGFP knocked into the *Scn2a* locus. Note GFP expression in granule cell layer and molecular layer and lack of expressing in Purkinje cell layer. **C:** Coronal section through cerebellar vermis, perpendicular to major axis of Purkinje cell dendrites. Note periodicity of staining in molecular layer, consistent with lack of expression in Purkinje cell dendrites. **D:** Left: Two EPSCs (10 Hz) before and after 166 Hz burst LTP induction in *Scn2a*^*+/+*^ (black, n = 6 cells from 6 mice) and *Scn2a*^*+/-*^ (cyan, n = 10 cells from 4 mice). Inset, paired pulse ratio before and after LTP induction. *: p < 0.01, Wilcoxon signed-rank test. Right, first EPSC amplitude, normalized to pre-induction average. Data are binned each minute. Circles and bars are mean ± SEM. **E:** Left: APs generated by current injection (10-50 pA, 300 ms) in *Scn2a*^*+/+*^ (black) and *Scn2a*^*+/-*^ (cyan). Middle: APs (spikes) per 300 ms stimulation epoch for each current amplitude (top; lines and shadow are mean ± SEM of population, *: p < 0.05 of slope between 10-40 pA), and near-rheobase APs plotted as dV/dt vs voltage (phase-plane plot) from *Scn2a*^*+/+*^ and *Scn2a*^*+/-*^. Right: AP threshold (top) and peak dV/dt (bottom) in *Scn2a*^*+/+*^ (13 cells from 3 mice) and *Scn2a*^*+/-*^ (16 cells from 3 mice). *: p < 0.05, Mann Whitney test. Circles are single cells, boxes are medians and quartiles with 90% tails. **F:** Left to right: spontaneous Purkinje cell AP train in *Scn2a*^*+/+*^ and *Scn2a*^*+/-*^, single AP, phase-plane plot of all APs in train, and summary AP threshold and peak dV/dt in *Scn2a*^*+/+*^ and *Scn2a*^*+/-*^. **G:** Left, Parallel fiber volleys evoked in trains of 10 APs at 50 Hz or 20 APs at 166 Hz. Right, Fiber volley amplitude, normalized per volley to first event in train, then averaged across recordings. Lines and bars are mean ± SEM. 50 Hz (top) in *Scn2a*^*+/+*^ (black, n = 5 from 2 mice) and *Scn2a*^*+/-*^ (cyan, n = 7 from 4 mice). 166 Hz (bottom) in *Scn2a*^*+/+*^ (black, n = 9 from 3 mice) and *Scn2a*^*+/-*^ (cyan, n = 9 from 4 mice). **H:** Left, parallel fiber evoked EPSCs in Purkinje cells from train of 20 stimuli at 166 Hz. Right, charge transfer, normalized to transfer from EPSC1 in *Scn2a*^*+/+*^ (black, n = 10 cells from 2 mice) and *Scn2a*^*+/-*^ (cyan, n = 9 cells from 2 mice) conditions.

To determine the underlying cellular excitability deficits that impair plasticity, we examined intrinsic and synaptic excitability features of the granule cell-Purkinje cell circuit. Consistent with immunostaining localization of Na_V_1.2 (*31*), intrinsic excitability was impaired in granule cells in *Scn2a*^*+/-*^ mice. Granule cell APs were slower to rise, and fewer Aps were evoked per given somatic current stimulus (Fig. 3E). By contrast, APs in Purkinje cells, which lack Na_V_1.2 channels (*29, 37*), were unaffected by *Scn2a* heterozygosity (Fig. 3B, C, E). This suggests that changes to Purkinje cell firing rate plasticity *in vivo* are due to alterations in presynaptic granule cell input.

Granule cells often fire in bursts *in vivo*, with average instantaneous frequencies of 160-170 Hz (*38, 39*). These high-frequency bursts are critical for gain-down plasticity, as they are a major signal that evokes LTP that is impaired in *Scn2a*^*+/-*^ mice (*40*–*43*). A high axonal Na_V_ density is required to sustain a burst of APs at high-frequency (*44*), suggesting that lower Na_V_ 1.2 density in parallel fiber axons may impair transmission. To test this, we assayed AP conduction fidelity in parallel fibers. At a lower frequency (50 Hz), parallel fiber volleys were sustained at normal levels in *Scn2a*^*+/-*^ mice; however, fiber volley amplitude was attenuated markedly at the higher frequency used for LTP induction (166 Hz) (Fig. 3G). Since the amplitude of axonal APs affects transmitter release probability (*45*), these smaller fiber volleys would be expected to evoke less transmitter at granule-Purkinje synapses. To test this, we again stimulated parallel fibers at high-frequency and now recorded resultant EPSCs in Purkinje cells. Similar to fiber volley measurements, we found that EPSC charge transfer was impaired in *Scn2a*^*+/-*^ mice, with more pronounced deficits occurring later in trains when volley waveform was more markedly reduced (Fig. 3H). Together, these data suggest that deficits in VOR plasticity in *Scn2a*^*+/-*^ mIce are strongly associated with deficits in transmission required for LTP at parallel fiber to Purkinje cell synapses.

### *Scn2a* heterozygosity in granule cells alone is sufficient to alter VOR gain

*SCN2A* is expressed across the brain. However, our results suggest that both synaptic and behavioral impact in *Scn2a*^*+/-*^ mice are likely mediated entirely by *Scn2a* heterozygosity in granule cells alone. To test this, we crossed animals heterozygous for a conditional *Scn2a* knockout allele (*Scn2a*^*+/fl*^) to the alpha6-Cre driver line, which expresses Cre largely in cerebellar granule cells16,46. As in constitutive *Scn2a* heterozygotes, VOR gain was elevated at baseline and could not be reduced by gain-down induction protocols (Fig. 4A-C). Furthermore, Purkinje cell firing rate was not increased during or after VOR gain-down induction (Fig. 4E-G), confirming that heterozygous loss of Na_V_1.2 in granule cells alone can impair VOR gain-down plasticity.

**Figure 4:**
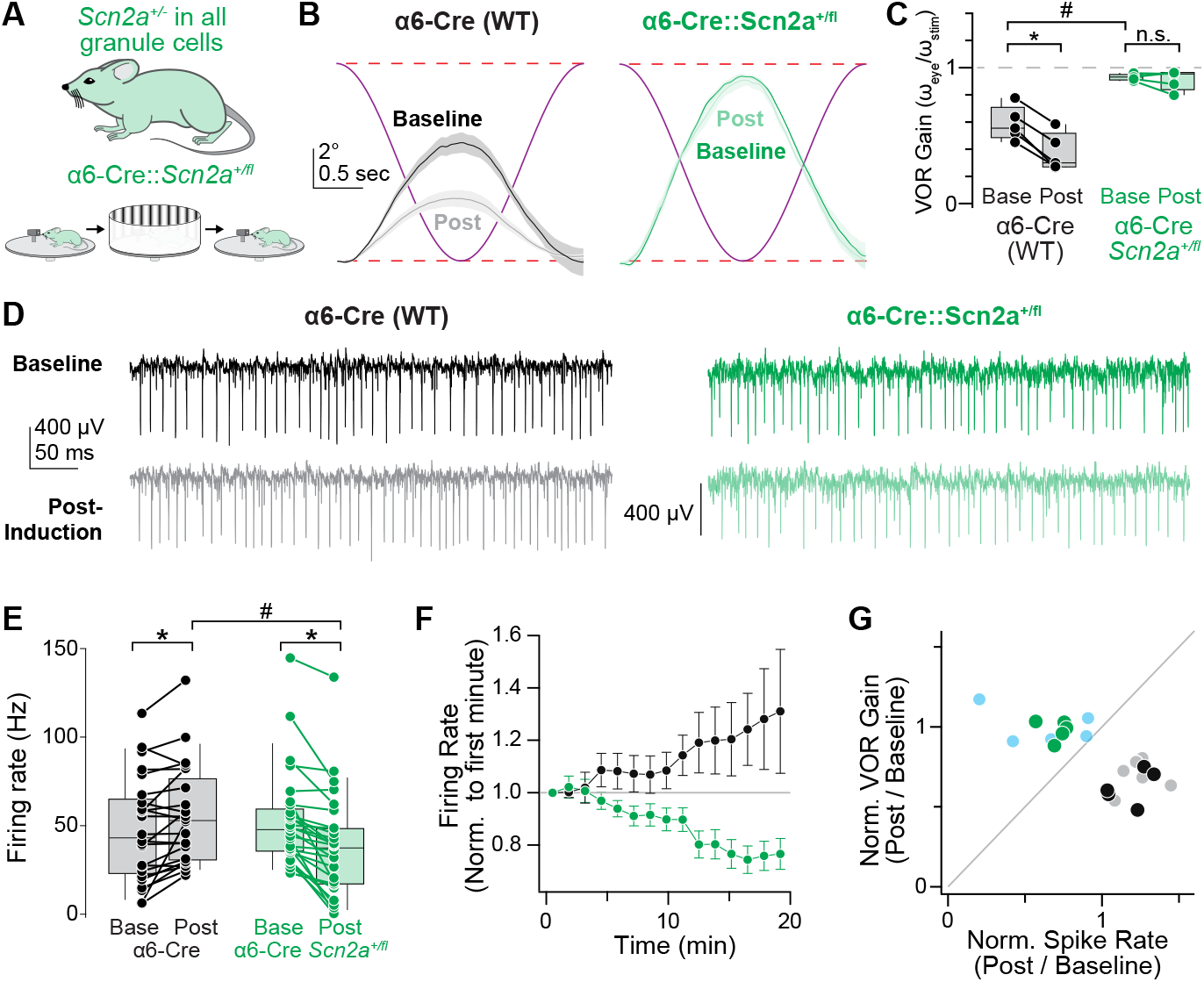
*Scn2a* heterozygosity in granule cells alone impairs VOR gain-down plasticity. **A:** Experimental design for VOR gain-down induction as in Fig. 2, but for *Scn2a*^*+/fl*^ mice crossed to the alpha6-Cre driver line (α6-Cre), which restricts Cre expression largely to cerebellar granule cells. **B:** Head angle (purple) and contraversive eye angle before (darker shade) and after (lighter shade) gain-down induction for α6-Cre not crossed to *Scn2a*^*+/fl*^ animals (black, WT-equivalent), or α6-Cre::*Scn2a*^*+/fl*^ mice (green). **C:** VOR gain before and after gain-down induction. Data color coded as in B. Bars connect data from individual mice. *: p < 0.001, Wilcoxon signed rank test, # p < 0.0001, Mann Whitney test. **D:** Putative Purkinje cell unit recordings before and after gain-down induction. Traces aligned to table rotation onset. **E:** Average Purkinje cell simple spike firing frequency during sinusoidal head rotation, before and after gain-down induction, displayed as in C. α6-Cre WT (gray, open circles, n = 24 units, 5 mice) and α6-Cre::*Scn2a*^*+/fl*^ mice (green, n = 28 units, 5 mice). *: p < 0.005, all conditions, Mixed-effects modeling, #: p = 0.01, Mann Whitney test. **F:** Average Purkinje cell simple spike firing frequency during induction protocol, normalized to firing rate in first minute per cell. Circles and bars are mean ± SEM, binned per minute. **G:** Normalized change in VOR gain vs. normalized change in simple spike rate (normalized per unit and averaged across units per animal). Circles are color-coded as in C and represent individual animals. Data from *Scn2a*^*+/+*^ (gray) and *Scn2a*^*+/-*^ (light cyan) are in background for comparison.

While the alpha6-Cre line allows for selective expression of Cre in granule cells within the cerebellum, there is some expression of Cre in other brain regions (*46*). To control for this, and to further confirm that floccular complex granule cell function is critical for VOR gain-down plasticity, we injected Cre bilaterally into the floccular complex of *Scn2a*^*+/fl*^ mice. Injections were performed at postnatal day 30 to further determine which components of VOR gain and its plasticity require normal Na_V_1.2 levels at this later state of development. Since granule cells are the only cell class at the injection site that express *Scn2a*, this approach will result in selective *Scn2a* heterozygosity in floccular granule cells alone. As observed in alpha6-Cre::*Scn2a*^*+/fl*^ mice, VOR gain-down plasticity was blocked. Interestingly, it did not alter baseline VOR gain from WT-like levels (Fig. S5), even 3 months after injection (Fig. S6). This suggests that baseline gain is maintained despite loss of gain-down plasticity, perhaps because there is no pressure to change baseline values once established in early development, as there is likely limited error in visual scene stabilization that would drive such changes. Consistent with this, VOR measured while presenting a static visual stimulus was near unity for both WT and *Scn2a*^*+/-*^ mice, confirming that *Scn2a*^*+/-*^ mice can respond to visual stimuli (Fig. S2).

### VOR gain-down plasticity can be rescued by upregulating *Scn2a* expression

These observations add to a growing body of literature showing that *Scn2a* is a key gene for synaptic and behavioral plasticity (*16, 47*–*49*). In neocortical circuits, we found recently that plasticity could be reinvigorated if *Scn2a* expression was restored in *Scn2a*^*+/-*^ mice (*50*). This was achieved using a CRISPRa-based therapeutic approach that targets a transcriptional activator to the residual, functional allele present in *Scn2a* heterozygotes. Since neurons are non-dividing cells, a single CRISPRa administration can effectively upregulate gene activity throughout life. Furthermore, previous research has shown that this approach only upregulates genes in tissues where the targeted regulatory element is active (*51*). Therefore, *Scn2a* upregulation would only occur in cells that express *Scn2a* endogenously, such as cerebellar granule cells. To test whether cerebellar plasticity could be similarly rescued with this approach, we injected a pair of blood-brain-barrier penetrant adeno-associated viruses containing all necessary CRISPRa components and the fluorescent protein mCherry via the retro-orbital sinus at P30 to upregulate *Scn2a* across the brain. To assess viral infection levels across cerebellum, we performed quantitative PCR (qPCR) for *Scn2a* mRNA in cerebellar lysates from all CRISPRa injected animals. The qPCR results showed elevation in normalized *Scn2a* mRNA expression levels in 5 of the 6 injected animals (Fig. 5A-C). Remarkably, VOR gain-down plasticity was restored in those 5 animals, but not in the last animal (Fig. 5D). Plasticity restoration coincided with increases in Purkinje cell firing rate during and after VOR gain-down induction (Fig. 5E-G), suggesting that both synaptic and behavioral plasticity could be rescued even when restored in later stages of development.

**Figure 5:**
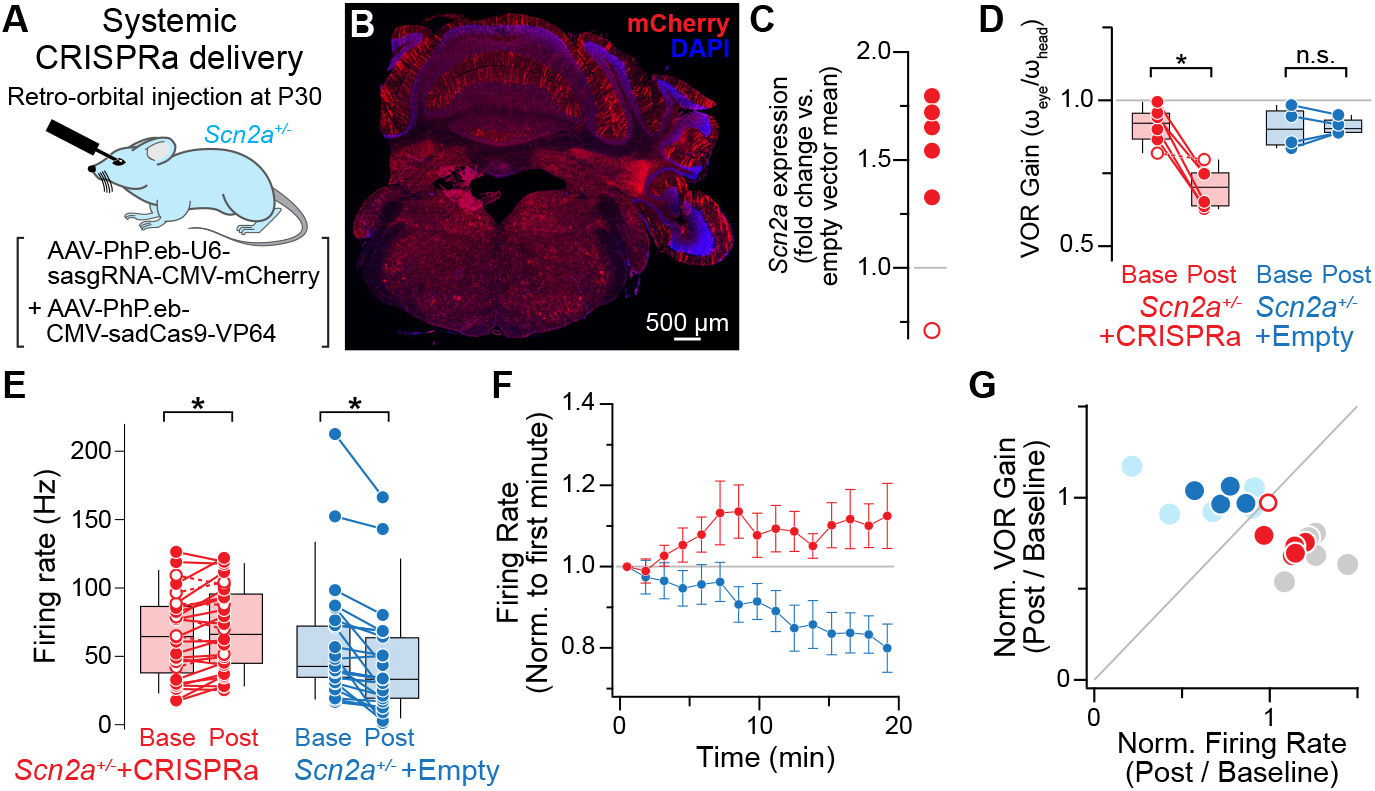
Rescue of VOR gain-down behavioral plasticity with CRISPR activation. **A:** Two AAV-PHP vectors are injected together for systemic infection via the retro-orbital sinus. AAV1 contains a guide RNA and mCherry. AAV2 contains dCas9 and the transcriptional regulator VP64. **B:** Cerebellar coronal section from injected animal. Note broad, but incomplete, infection across cerebellum, as signaled by mCherry fluorescence. A region of cerebellum containing the left floccular complex was dissected immediately after harvesting brain for quantitative PCR and is missing from section. **C**. Quantitative PCR result showing *Scn2a* mRNA detection in mouse cerebellum of “empty vector” controls, which contain all viral elements except the guide RNA. In 5/6 animals, marked upregulation was noted (closed circles). In 1 animal, no upregulation was noted (open circle). **D:** VOR gain before and after gain-down induction in *Scn2a*^*+/-*^ mice with *Scn2a* CRISPRa (red, 6 mice) and empty vector (blue, 4 mice), depicted as in Fig 2D. *: p < 0.05 Wilcoxon-signed rank test; all data included. **E:** Average Purkinje cell simple spike firing frequency during sinusoidal head rotation, before and after gain-down induction in *Scn2a*^*+/-*^ mice with CRISPRa (red, n = 28 units from 6 mice) and empty vector (blue, n = 25 units from 4 mice). Circles are color-coded as in D and represent individual units. *: p < 0.005, Mixed-effects modeling. **F:** Normalized Purkinje cell simple spike firing frequency during gain-down induction for all CRISPRa (6 mice) and empty vector (4 mice) injected animals. Circles and bars are mean ± SEM, binned per minute. **G:** Normalized change in VOR gain vs. normalized change in simple spike rate (normalized per unit and averaged across units per animal). Circles are color-coded as in D and represent individual units. Data from *Scn2a*^*+/+*^ (gray) and *Scn2a*^*+/-*^ (light cyan) are in background for comparison. Note overlap of CRISPRa population with *Scn2a*^*+/-*^ population (except open circle) and empty vector with *Scn2a*^*+/-*^ population.

## DISCUSSION

Oculo-motor reflexes are commonly assessed for diagnosis of neurological disorders. Here, we highlight how examination of VOR can shed light on dysfunction in cellular, circuit, and behavioral plasticity in neurodevelopmental disorders as well. VOR function is conserved across vertebrates. We observed that its dysfunction is also conserved in mouse and human *SCN2A* haploinsufficiency. This parallels observations in non-syndromic ASD, where heightened VOR gain or dysfunction in other aspects of vestibular sensation were also observed (*10, 12*). VOR is detectable within the first months of life in humans (*52, 53*), well before ASD is typically diagnosed (*54*). Given the importance of early diagnosis (*55*), examination of VOR and related innate behaviors may help improve patient and family outcomes. Furthermore, we show that plasticity in VOR gain is sensitive to therapeutic intervention in mouse models.

Here, deficits in VOR were due specifically to deficits in granule cell transmission. This, in turn, altered mechanisms for regulation of Purkinje cell-mediated inhibition of VOR circuit through the vestibular nuclei. The net result, for both human and mouse models of *SCN2A* LoF, was a VOR gain saturated near unity, even at slow head angular velocities. These effects appear due to alterations at parallel fiber-to-Purkinje cell synapses. Though granule cells also synapse onto molecular layer interneurons that provide feedforward inhibition to PCs, interneuron activity is not necessary for gain-down plasticity (*33*). Furthermore, our understanding of interactions between parallel fiber firing rates and plasticity at synapses onto Purkinje cells is consistent with impaired LTP in *Scn2a*^*+/-*^ conditions. At parallel fiber-to-Purkinje cell synapses, high-frequency bursts of APs are required for LTP related to VOR gain-down plasticity (*34*). By contrast, long-term depression can be supported by shorter bursts of granule cell activity paired with Purkinje cell depolarization, and likely remains intact even in *Scn2a*^*+/-*^ conditions (Fig. 3F) (*42, 56*–*58*). Thus, loss of LTP may bias VOR gain towards unity and render VOR inflexible to gain-down protocols.

The simplicity of cerebellar anatomy — composed of relatively few cell classes with stereotyped synapses — belies its functional complexity. Indeed, the cerebellum contributes to a range of processes, including social and cognitive processing that are affected in ASD (*59*–*64*). Here, we built upon decades of work that established a role for parallel fiber-Purkinje cells synaptic plasticity and VOR adaptation to understand links between cellular dysfunction and alterations in reflexive behavior in *SCN2A* haploinsufficiency. While this behavior requires the floccular complex alone, NaR_V_1.2 is expressed ubiquitously in granule cells across the cerebellum, as is high-frequency, burst-based LTP to Purkinje cells (*40, 41, 43, 65*). Thus, vestibular plasticity deficits observed here may hint toward broader cerebellar plasticity deficits, affecting a range of phenotypes in children with *SCN2A* LoF (*15*). Furthermore, this work parallels studies of cerebellar plasticity in mouse models of other ASD-associated genes (*66*–*70*). Thus, cerebellum-dependent reflexes may prove to be generalizable and translatable biomarkers for ASD and potential therapeutics, allowing for quantitative assessment of behavior throughout development.

### Limitations of the study

This work represents the first study, to our knowledge, in which VOR gain was assessed in children with profound developmental delay. Children with *SCN2A* LoF, including those in this study, are often non-verbal, unlikely to follow complex instruction, and, from consultation with caregivers, very unwilling to undergo head restraint in apparatus traditionally used to assess VOR gain in the clinic. Given these limitations, we designed a helmet-mounted VOR apparatus that allows a child to move more freely and restricted our analysis to baseline VOR gain. In contrast to our experiments in *Scn2ar*^*+/-*^ mice, we did not attempt to assess VOR gain plasticity with any induction protocols in children. In future studies, this may be possible, perhaps by using virtual reality headsets.

This study was limited to children that present with *SCN2A* LoF without seizure or anti-seizure medication within the past 6 months. An estimated 20-30% of children with LoF variants present with seizure, usually with an onset after the first 3-6 months of life (15). Seizures are treated with a range of medications, but those that interfere with sodium channel function are typically avoided (71). How such medications interact with the physiology that underlies VOR is unknown. Furthermore, mouse models with constitutive or conditional *Scn2a* deletion were used. While this models the majority of LoF cases in the human population, it does not model missense variants that either dampen or eliminate channel function, but do not lead to nonsense mediated decay of NaR_V_1.2. How VOR is encoded in such conditions will require creation of new missense models.

## Supplementary materials

Supplementary materials contain Methods and Supplemental Figures 1-6.

## Acknowledgments

We are grateful to the children, parents, and caregivers associated with the FamilieSCN2A Foundation for participating in this study. We thank Drs. B. Barbour, M. Casado, J. Christie, F. Dunn, A. Nelson, P. Reeson, M. Scanziani, and M. Schonewille for feedback on this work, and to Dr. J. Christie for discussions that motivated this study. This work was supported by an Action Potential Award from the FamilieSCN2A Foundation (CW), a research grant from BioMarin Pharmaceutical Inc. (KJB), the Weill Neurohub Investigator Program (KJB, NA), SFARI 629287 (KJB, NA), Howard Hughes Medical Institute funds awarded to M. Scanziani (GB), and National Institutes of Health grants F32 MH125536 (ADN) and R01 MH125978 (KJB).

## Author contributions

Conceptualization—CW, GB, KJB

Methodology—CW, KDD, EH, XZ, ADN, HK, NA, GB, KJB

Software—CW, KDD, HK, GB Formal analysis—CW

Investigation— CW, KDD, EH, XZ, ADN, HK, NA, GB, KJB

Writing - Original Draft—CW

Writing - Review & Editing—All authors Visualization—CW, GB, KJB

Supervision—NA, GB, KJB

Funding acquisition—CW, ADN, NA, KJB

