## Supplemental Material for "Impaired cerebellar plasticity hypersensitizes sensory reflexes in *SCN2A*-associated ASD"

### Supplemental Figures 1-6

### Methods

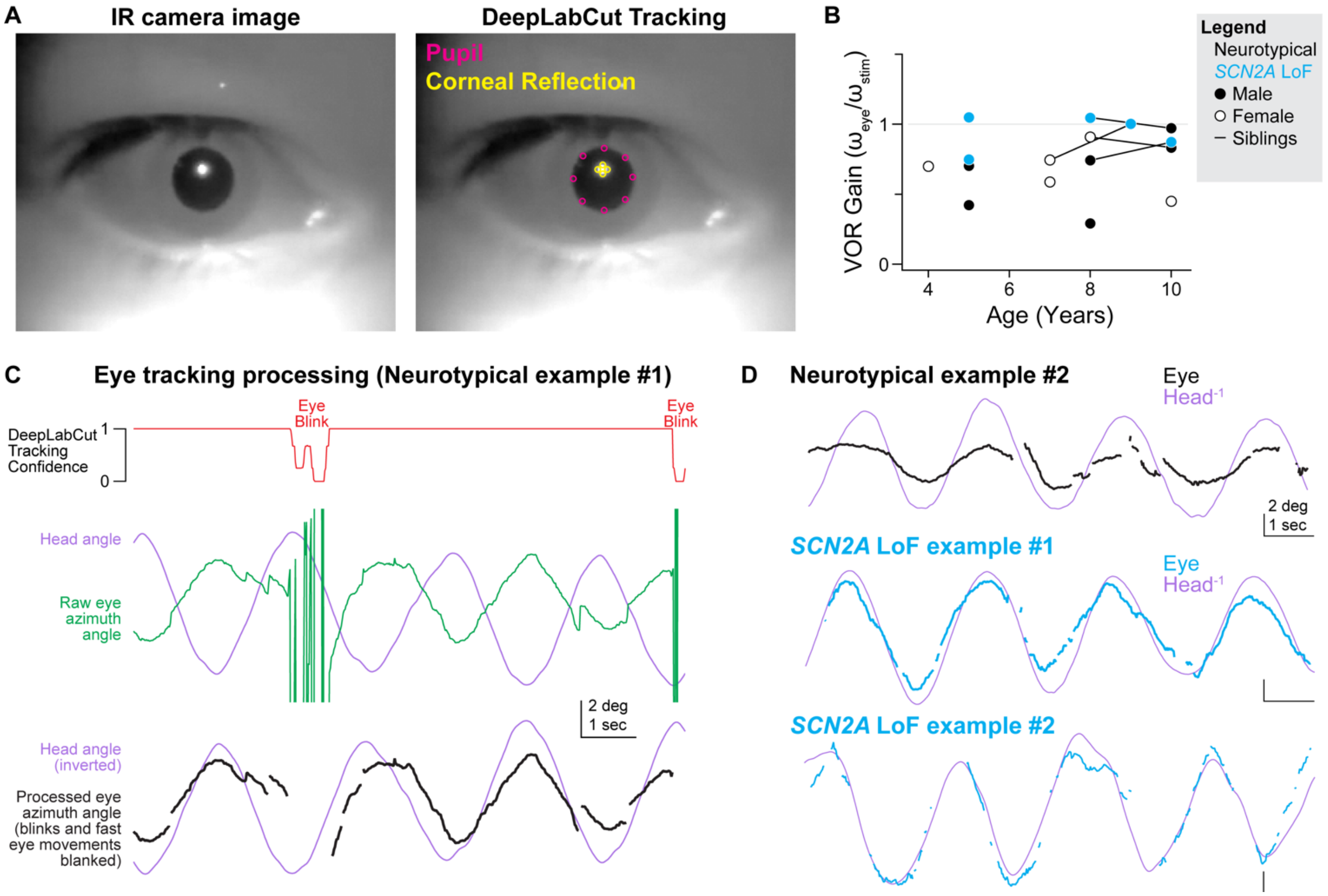


**Figure S1: Human VOR analysis**

A: Example infrared image of right eye (left) and overlaid DeepLabCut tracking of pupil boundaries and corneal reflection (right). Landmarks were tracked at 8 and 4 locations near areas of maximal contrast to minimize tracking errors. Azimuth position was calculated by averaging across all points.

B: VOR gain per subject across age, coded by sex and *SCN2A* phenotype. Siblings are connected by lines.

C: Analysis pipeline for VOR quantification. Top: Red trace is average DeepLabCut confidence metric of marker position. Middle: purple trace is head angle determined from inertial motion unit. Green trace is raw DeepLabCut eye angle. Bottom: Black trace shows post-processing of eye position after periods of low DeepLabCut confidence and periods where eyes move >20°/s are blanked. Note that the head never moved faster than 12.5°/s during 0.4 Hz rotations.

D: Other examples of post-processed eye and head angle in neurotypical and *SCN2A* LoF children.

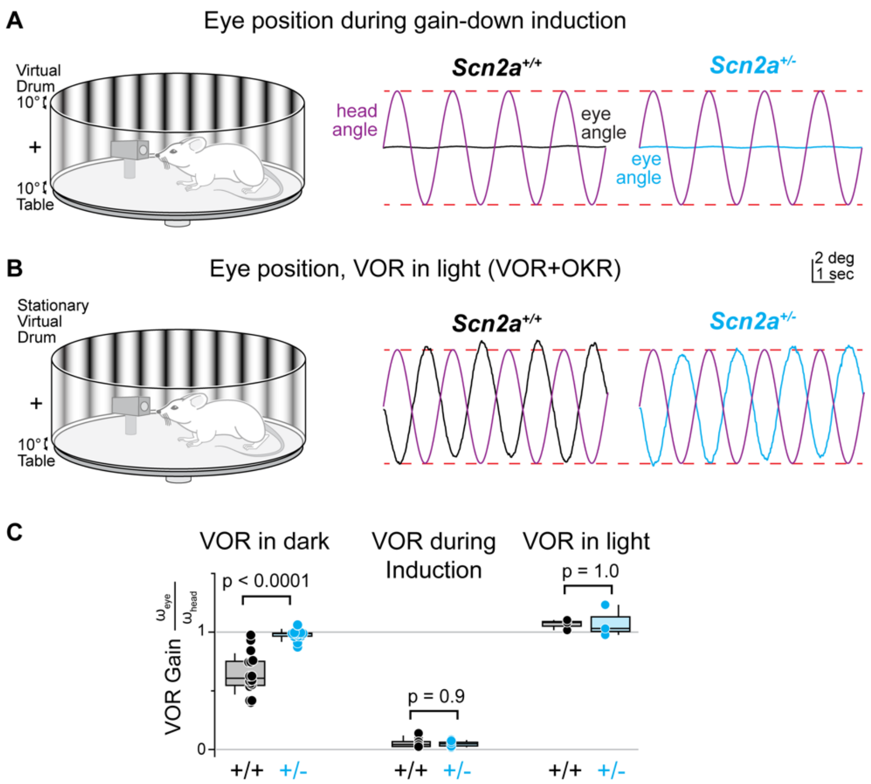


**Figure S2: Eye movement during gain-down induction and VOR in light**

A: Left: schematic of experimental setup for VOR gain-down induction. Right: example of eye position during VOR gain-down induction in WT and *Scn2a*$^{+/-}$ mice.

B: Left: schematic of experimental setup for VOR in light, where the mouse is rotated while the visual stimulus remains fixed. Right, example eye position in WT and *Scn2a*$^{+/-}$ mice.

C: Data summarizing gain during VOR in the dark (left), during gain-down induction (middle), and VOR in light (right). Circles are animals, boxes are medians and quartiles with 90% tails. P-values are for Mann-Whitney comparisons.

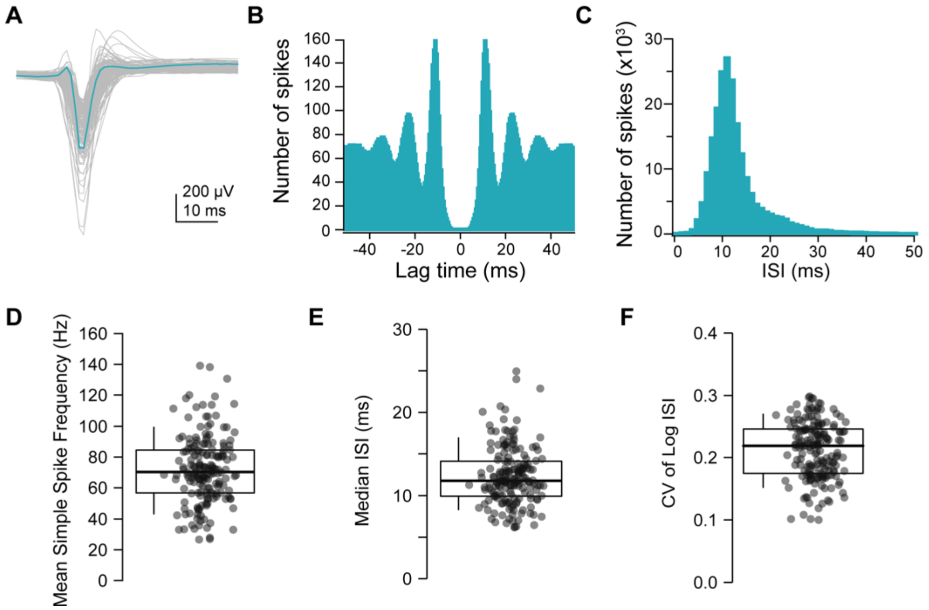


**Figure S3: Properties of putative Purkinje cell simple spike units**

A: Overlayed median simple spikes waveform from each putative Purkinje cell unit in all mice (144 units from 39 animals).

B: Autocorrelogram of the example Purkinje cell unit in A during the entire recording session.

C: Inter-spike interval distribution of the example Purkinje cell unit in A.

D: Distribution of mean simple spike firing frequency from all included Purkinje cell units.

E: Distribution of median inter-spike interval (ISI) from all included Purkinje cell units.

F: Distribution of the coefficient of variation of the natural log of the ISIs from all included Purkinje cell units.
In each panel, circles are units. Boxes are median and quartiles, with 90% tails.

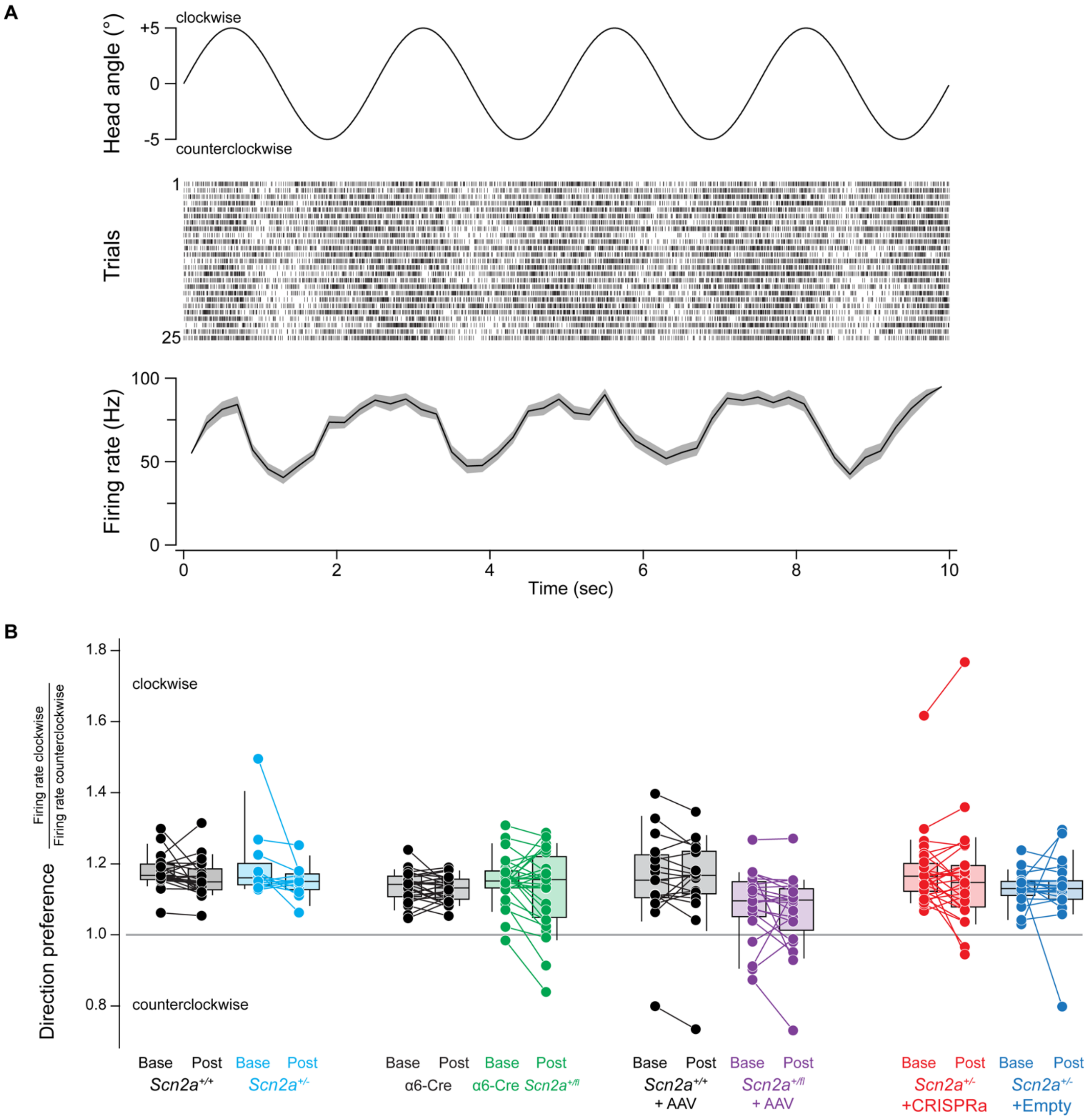


**Figure S4: Putative Purkinje simple spiking units display directional bias to head rotation**

A: Example Purkinje cell unit activity during table rotation (VOR baseline, WT animal). Top is head angle. Middle is simple spike raster across all 25 trials. Bottom is mean ± SEM firing rate, binned every 200 ms.

B: Summary direction preference (total spikes during clockwise movement / total spikes during counterclockwise movement) for each unit across all conditions. Circles are single units; lines connect individual units. Box plots are medians and quartiles with 90% tails. Note that the majority of units prefer clockwise movement.

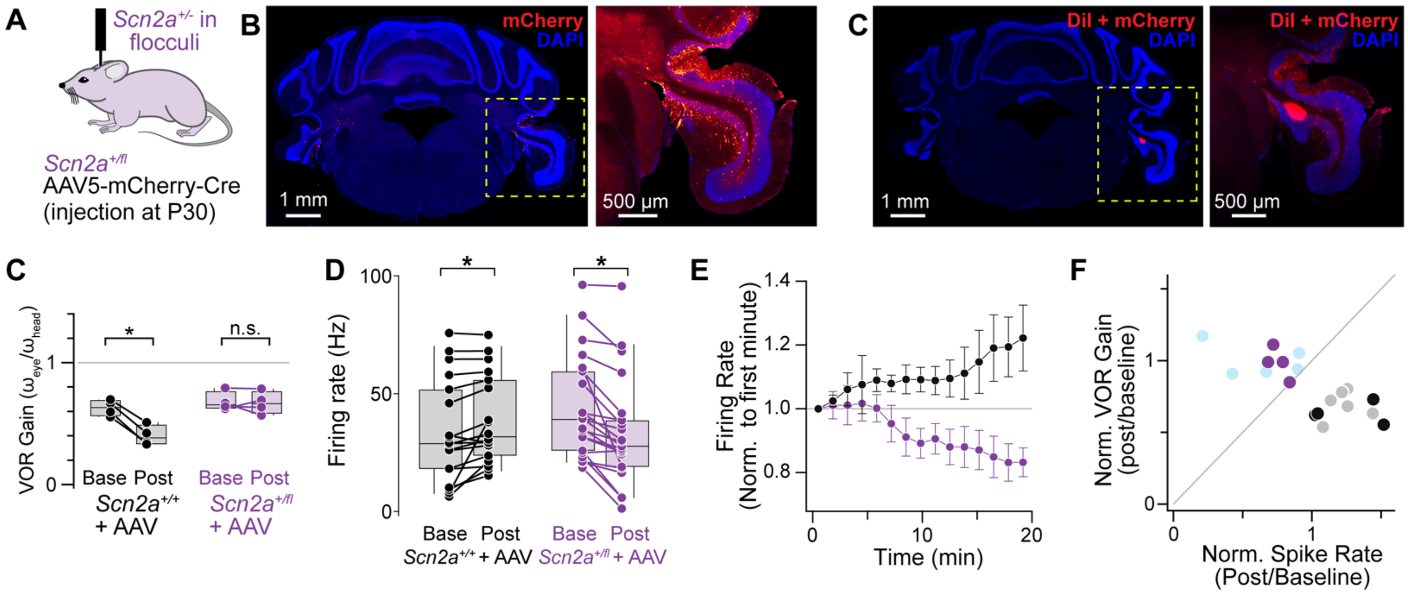


**Figure S5: *Scn2a* heterozygosity induced only in the floccular complex impairs VOR behavioral plasticity**

A: AAV5-mCherry-Cre was injected bilaterally into the floccular complex in *Scn2a*$^{+/fl}$ mice at P30.

B: Coronal section through floccular complex detailing mCherry expression. Yellow dashed box is expanded on right, using the "red-hot" lookup table (ImageJ) for mCherry fluorescence.

C: Coronal section detailing identification of silicon probe with DiI in animal previously injected with AAV5-mCherry-Cre.

D: VOR gain before and after gain-down induction in *Scn2a*$^{+/+}$ mice with Cre (black, N = 4 mice) and *Scn2a*$^{+/fl}$ with Cre (purple, N = 4 mice). *: p = 0.029, Wilcoxon signed rank test.

E: Average Purkinje cell simple spike firing frequency during sinusoidal head rotation, before and after gain-down induction in *Scn2a*$^{+/+}$ with Cre (red, n = 18 units from 4 mice) and *Scn2a*$^{+/fl}$ with Cre (purple, n = 21 units from 4 mice). *: $p < 0.05$, Mixed-effects modeling.

F: Normalized Purkinje cell simple spike firing frequency during gain-down induction. Data are binned every minute. Circles and bars are mean ± SEM.

G: Normalized VOR gain vs normalized simple spike firing frequency following gain-down induction for each mouse. *Scn2a*$^{+/+}$ (gray) and *Scn2a*$^{+/-}$ (light blue) data are displayed for comparison.

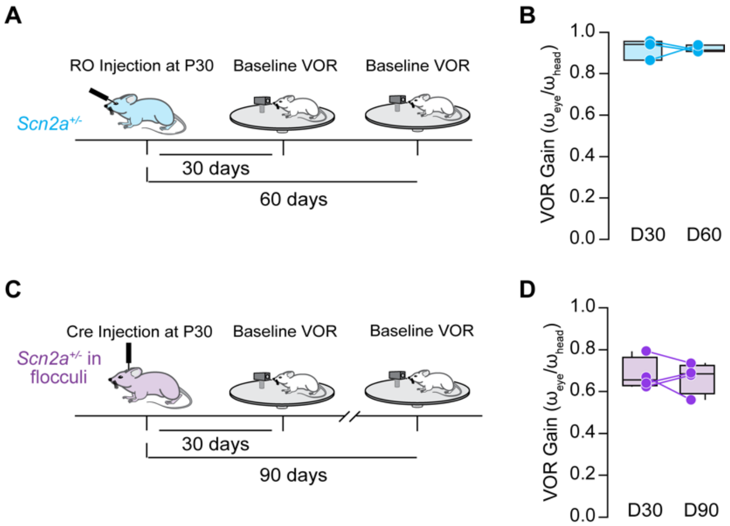


**Figure S6: Baseline VOR gain remains unchanged after manipulations of *Scn2a* expression from P30.**

A: Timeline of CRISPRa RO injection (at P30) and VOR recordings (30 days and 60 days after injection).

B: Data summarizing baseline VOR gain at post-injection day 30 and day 60 in *Scn2a*$^{+/-}$ mice with *Scn2a* CRISPRa.

C: Timeline of Cre injection (at P30) and VOR recordings (30 days and 90 days after injection).

D: Data summarizing baseline VOR gain at post-injection day 30 and day 90 in *Scn2a*$^{+/fl}$ mice with Cre.

**Methods:**

**Human vestibulo-ocular reflex:**

Experiments were performed on children of either sex, aged 3-10 years. All procedures were in accordance with UCSF IRB guidelines and parental consent was obtained for all participants. Inclusion criteria for *SCN2A* loss-of-function (LoF) children included those with a genetic diagnosis of *SCN2A* dysfunction and a diagnosis of autism spectrum disorder and neurodevelopmental delay. Furthermore, children were limited to those without seizure or any seizure-related medication within the last 6 months. Neurotypical (Nt) children were limited to those that did not present with neurodevelopmental delay, autism spectrum disorder, or seizure.

For VOR experiments, children were seated in a standard office executive chair with the center of their head situated near the azimuth swivel. Children were either seated alone or in a caregiver's lap for both Nt and LoF cohorts, with no discernable differences in data quality due to caregiver presence. Children were then fitted with a helmet (67i skateboard helmet, child's small or medium depending on participant, Amazon) that was kitted with an inertial movement unit centered at the top of the helmet and an infrared camera (Raspberry Pi NoIR) and 940 nm LED (Luxeon star) with a 10° diffuser lens focused on the right eye. The camera was aligned to ensure that a corneal reflection from the LED could be visualized. Eye movement was first calibrated in light conditions by having children attend to a visual stimulus (a chirping yellow sparrow stuffed animal or a pen light clicked on and off as conditions permitted) moved back and forth from a position either directly ahead or 10 degrees offset. Calibration was repeated >4x for each child and eye position calibration tracking was averaged over all trials.

Following calibration, the chair was rocked back and forth ~±5° at ~0.4 Hz in the dark by an experimenter wearing night-vision optics, with oscillations paced by a metronome. Videography and IMU data were acquired at 60 and 300 Hz, respectively. IMU data were downsampled to 60 Hz for subsequent analysis.

Data acquisition could not be performed blind to LoF/Nt condition given the phenotypes being analyzed. All data was therefore coded and blinded for subsequent analysis. Post-hoc processing of eye position was performed using DeepLabCut 2.0 Toolbox (deeplabcut.org) (*1*). DeepLabCut was trained by an experimenter to detect 8 points at regions of maximal contrast at the edge of the pupil (every 45 degrees) and 4 points at the edge of the corneal reflection (every 90 degrees). Horizontal eye angle was calculated as the average horizontal pupil value (all 8 points) relative to the average horizontal corneal reflection value (all 4 points). Data acquired during VOR tests were then normalized to calibration sets to obtain the exact eye movement angle.

All children naturally made saccades and blinked during data acquisition. These events were removed from subsequent analysis by eliminating all data when eye movement speed exceeded 20 deg/sec and where the DeepLabCut position "confidence" value, averaged over all 12 imaged points, was <0.99 (range: 0 to 1.0). Eye angle and head angular velocity (IMU) data were used to calculate instantaneous velocity, smoothed with a 2 data-point-wide Gaussian filter. VOR gain was then calculated as the ratio of the integral of absolute values of instantaneous eye velocity and corresponding values of head velocity over the entire recording epoch.

**Animals:**

All experimental procedures were performed in accordance with UCSF IACUC guidelines. All experiments were performed on mice housed under standard conditions with *ad libitum* access to food and water, with colonies maintained in-house. Wild-type C57B6J were from JAX (000664); Alpha6-Cre mice were initially described in Bahn et al., 1997 (*2*); *Scn2a*$^{+/-}$ mice were initially described in Plannels-Cases, 2000 (*3*); conditional knockout *Scn2a* allele mice were initially described in Spratt et al., 2019 (*4*); conditional knockin *Scn2a* mice with GFP in-frame were described in Tamura et al., 2022 (*5*).

**Surgical procedures:**

*Head Plate Implantation for Head-fixed Recordings*

Mice were implanted with a head bar composed of three threaded screw inserts arranged in a triangle (McMaster-Carr, 92395A109) at least 14 days before recording. Mice were anesthetized with 2% isoflurane. The scalp was shaved and disinfected with povidone iodine. The scalp was resected, and the skull was scored with a sterilized bone scraper. The edge of the skin was glued to the skull and the metal threaded screw inserts were sterilized and mounted using Metabond and dental cement (Ortho-Jet powder; Lang Dental) mixed with black paint (iron oxide). The inserts were mounted via a stereotax with the triangular center aligned to bregma. Following the implantation, black dental cement was used to build a recording well surrounding the skull above the targeted floccular complex region at (from the bregma, in mm): anterior-posterior, -5.40, mediolateral, -2.75. The surface of the skull above the left hemisphere of the cerebellum was not covered with dental cement but was coated with a thin layer of transparent cyanoacrylate glue. Mice were injected intraperitoneally with 0.1 mg/kg buprenorphine before and after surgery and were checked daily for 3 days after the head-bar surgery.

*Craniotomy for Electrophysiological Recordings*

Following the last day of habituation to head fixation, each mouse was anesthetized with 2% isoflurane and the skull above the recording site was drilled off. The dura was not removed, and the craniotomy was kept moist with phosphate buffered saline (PBS). Following craniotomy, the window was sealed with biocompatible silicone sealant until the recording session (Kwik-Cast, World Precision Instruments).

*Stereotaxic transcranial viral injection*

Mice with one conditional knockout *Scn2a* (*Scn2a*$^{+/fl}$) allele at P30 were anesthetized under isoflurane at 2.0% and mounted onto the stereotaxic machine (Kopf 1900). 500 nL of AAV-EF1a-mCherry-IRES-Cre-WPRE (UNC Vector Core) was injected into the floccular complexes on both sides at (from the bregma, in mm): anterior-posterior, -5.40; mediolateral, ±2.75; dorsoventral: −4.00 at 100 nL per minute.

*Viral injection via retro-orbital sinus*

AAV-PhP.eb-CMV-sadCas9-VP64 with either AAV-PhP.eb-U6-sasgRNA-CMV-mCherry (CRISPRa) or an empty vector containing AAV-PhP.eb-U6-CMV-mCherry suspended in 5% Sorbitol (Sigma-Aldrich, Inc.) at a total volume of 150 µL and packaged in a 28-gauge needle with 0.5 mL insulin syringe (U-100; Becton, Dickinson and Company). Mice aged P30 were anesthetized under inhalant isoflurane and were kept warm on a heating pad. Ophthalmic ointment (Puralube Vet Ointment; Dechra) was applied to both eyes of the anesthetized mouse. The mouse was then laid on its left side and the needle was subsequently inserted into the medial canthus underneath the left eyeball until it reached the base. The mixture of viral constructs was then introduced into the retro-orbital sinus. After a 10-second pause, the needle was slowly withdrawn and additional ocular ointment was applied. The mice were monitored for signs of bleeding (none noted) and provided analgesic and monitored for pain for 3 days post-injection (*6*).

**Behavior:**

*Habituation and eye calibration*

Behavioral experiments were performed during the light cycle and the experimenter was blind for genotype during experiment and analysis. After allowing for at least 1 week of recovery after surgery, each mouse was habituated to the experimental setup by head-fixing it in a purpose-built restraining tube with a head holder that was positioned at the center of the turntable for 1-2 hours each day for 4 consecutive days (*7*). Each mouse was habituated for 1 hour in the darkness on the first day, then with a 20-minute increment on each subsequent day for the following three days.

On the third habituation day, the right eye of each mouse was calibrated using a standard method, as reported previously (*7*). Briefly, to calibrate eye movement, we used a corneal reflection generated by an infrared LED as a reference position for the eye. A static virtual drum was presented during calibration. After identifying the pupil and the center of the eye, the camera and the reference LED were rotated by 10 degrees in both directions to acquire the distance between the pupil reference and the pupil center. The pupil size was varied by changing the luminance of the virtual drum to obtain increase the dynamic range of pupil size values.

*Vestibulo-ocular reflex recordings*

Mice were given 3% topical pilocarpine HCl in both eyes 15 minutes prior to the beginning of each VOR recording session to temporarily restrict their pupil size in the dark. Compensatory eye movements were recorded using video-oculography. The velocity of head movements was controlled by the servo-controlled platform that enabled the rotation of the animal along the horizontal plane. The platform was attached to a gearbox 15:1 (VTR010-015-RM-71 VTR, Thomson) that increased the torque of a servo motor (AKM53L-ANC2C-00 KEC0432 AC Servomotor 1.83 kW, Kolmorgen). The motor was tuned using a servo drive (AKDB013206-NBAN-0000 servo drive, Kolmorgen) and controlled in velocity mode using analog waveforms computed in LabView 2020 (NI). The head of each mouse was positioned with a 20 degrees angle along the sagittal plane of the head (nose pointing 20 degrees down), such that the plane of the horizontal vestibular canal was approximately parallel to the plane of the rotating platform. VOR was evoked by rotating the platform in the dark with 5° amplitude in each direction and a temporal frequency of 0.4 Hz unless otherwise stated. Gain-down VOR induction protocol was performed by presenting a coherent virtual drum with a spatial frequency of 0.8 Hz and a temporal frequency of 0.4 Hz with the platform rotation that oscillates in the same direction to evoke near complete VOR cancellation (see Figure S2A).

#### *In vivo* Extracellular recordings:

All recordings in this study were performed in the left floccular complex (contralateral to the imaged eye). Extracellular recordings were performed using a 2-shank silicon probe that is 8 mm in length and contains 32 channels on each shank with 25 µm inter-channel distance and 250 µm inter-shank distance (ASSY-77 H2, Cambridge Neurotech). The recording electrode was controlled with a micromanipulator (MP-285; Sutter Instrument) and stained with DiI lipophilic dye (Thermo Fisher) for post-hoc identification of the electrode location (see Figure S5C). The signals were acquired at 30 kHz using an INTAN system (RHD2000 USB Interface Board, INTAN Technologies). Recordings in the floccular complex were performed at (from bregma, in mm): anterior-posterior, -5.40; mediolateral, -2.75; dorsoventral: −4.00.

On the day of recording, each mouse was head-fixed on the VOR experiment setup. The silicon sealant was then removed, and the craniotomy was kept moist with artificial cerebrospinal fluid contained (in mM): 140 NaCl, 5 KCl, 10 D-glucose, 10 HEPES, 2 CaCl2, 2 MgSO4 with a pH of 7.4. The recording electrode was lowered into the left floccular complex via the craniotomy at a speed of ~250 µm per minute. Experiments began at least 50 minutes after reaching the final electrode depth to allow for electrode stabilization.

#### Histology:

For anatomical analysis, mice were perfused transcardially with phosphate buffered saline (PBS, Sigma-Aldrich) and then with 4% paraformaldehyde (PFA, VWR International, Inc.) in PBS. Brains were extracted from the skulls, post-fixed in 4% PFA overnight at 4°C, switched to 15% sucrose in PBS for 4 hours, and then stored in 30% sucrose in PBS until sectioned. Sequential coronal sections of the cerebellum (30 µm) were collected with a Microm HM 525 Cryostat and were stored in 0.05% sodium azide (Sigma-Aldrich) in PBS. For *in vivo* recording electrode location identification, sections were rinsed in 1X PBS 3 times for 15 minutes and then were then coverslipped with ProLong Gold with DAPI (Life Technologies). For immunohistochemistry mCherry staining, sections were rinsed in 1X PBS 3 times for 15 minutes and then incubated overnight at 4°C with 10% normal goat serum (Fisher Scientific) with 0.2% Triton X-100 (Sigma-Aldrich) and Anti-RFP (Rabbit) primary antibodies (1:1000) (Rockland Immunochemicals). The next morning, sections were kept at room temperature for 1 hour before being rinsed with 1X PBS 3 times for 15 minutes. The sections were then incubated with Alexa Fluor 568 goat anti-rabbit (1:500) (Invitrogen) for 2 hours at room temperature. Sections were then coverslipped with ProLong Gold with DAPI (Life Technologies). Fluorescence images were acquired using an Olympus FV3000 confocal microscope under 4x or 20x objectives.

#### Data Analysis:

*Monitoring eye movements by video-oculography*

The movement of the right eye was monitored using a high-speed infrared (IR) camera (IPX-VGA210LMCN; Imperx, Inc.) by capturing the reflection of the eye on an IR mirror (Edmund Optics #64–471) that is transparent to visible light under the control of custom routines in LabView 2020 (NI) and a frame grabber (PCIe-1427, NI) at an acquisition rate of 100 Hz. The pupil was identified online by thresholding pixel values and its profile was fitted with an ellipse to determine the center. The eye position was measured by computing the distance between the pupil center and the corneal reflection of a reference IR LED placed along the optical axis of the camera.

*VOR gain calculation*

Eye tracking data were processed in customized routines in MATLAB. VOR gain was calculated as the ratio of the average of the difference of the maximum and minimum of the eye position for each sinusoidal cycle and the amplitude of table rotation (10 degrees, 10s trial duration for all rotation frequencies). VOR gain was calculated using sinusoidal cycles that did not contain saccades. At least 20 trials were collected from each mouse.

*Purkinje cell simple spikes identification*

Automated spike sorting was carried out using KiloSort (https://github.com/cortex-lab/Kilosort) (*8*) by manual curation of the units using Phy2 (https://github.com/cortex-lab/phy). Multiple parameters were used to identify putative Purkinje cell units. First, we only included units that exhibited spikes across at least 3 adjacent channels on the linear probe (25 μm inter-channel distance) to bias selection towards cells that have large somatodendritic area. This is thought to exclude units arising from molecular layer interneurons (*9*). Second, we excluded units with refractory period violations greater than 1-2%. Simple spikes from putative Purkinje cell units were then validated using previously defined standards based on mean firing rates (> 22 Hz), median inter-spike intervals, inter-spike interval (ISI) distributions, and the coefficient of variation of the natural log of the ISIs (Range: 0.05 to 0.4) (*10*). Purkinje cell units outside of these ranges were excluded from subsequent data analysis. We also excluded complex spikes from our analysis.

*Firing frequency and CV of ISI computation*

All analysis was carried out via MATLAB (MathWorks) and Python on identified Purkinje cell units. The timings of each simple spike were aligned with individual trials of VOR recordings using a digital pulse triggered by a photodiode positioned on the virtual drum. Only simple spikes that occurred during the 10-second sinusoidal oscillation trials were included in subsequent analyses. To determine directional bias of the recorded PC units, we used the ratio of simple spikes frequency during clockwise (CW) and counterclockwise (CCW) head rotation in each trial. Overall simple spike firing frequency was calculated by treating each trial (10 seconds) as a bin and dividing the total number of spikes by the duration of the trial. The firing frequency from each trial was then averaged within each experimental epoch, including pre- and post-gain down induction. The firing frequency of each trial during gain-down induction was calculated in the same way but was binned every 4 trials and normalized to the first bin to show percentage change over time. The coefficient of variation (CV) of the inter-spike intervals (ISI) was calculated on the same set of Purkinje cell simple spikes.

### ***Ex vivo* Electrophysiology:**

*Ex vivo cerebellar tissue preparation*

Mice aged P28 through P59 were anesthetized and 250 μm-thick acute coronal slices containing the cerebellum were prepared. Slices were prepared from *Scn2a*$^{+/-}$ or wild-type littermates (genotyped by PCR). All data were acquired and analyzed blind to *Scn2a* genotype. Data were acquired from both sexes (blind to sex). Cutting solution contained (in mM): 87 NaCl, 25 $NaHCO_3$, 25 glucose, 75 sucrose, 2.5 KCl, 1.25 $NaH_2PO_4$, 0.5 $CaCl_2$ and 7 $MgCl_2$; bubbled with 5%$CO_2$/95%$O_2$; 4°C. Following cutting, slices were incubated in recording solution for 30 min at 33°C, then at room temperature until recording. Recordings solution contained (in mM): 125 NaCl, 2.5 KCl, 1.3 $CaCl_2$, 1 $MgCl_2$, 25 $NaHCO_3$, 1.25 $NaH_2PO_4$, 25 glucose; bubbled with 5%$CO_2$/95%$O_2$; 32-34°C, ~310 mOsm at ~33°C.

*Ex vivo electrophysiological recordings*

Cerebellar Purkinje cells and granule cells were visualized with differential interference contrast (DIC) optics for conventional visually guided whole-cell recording. For current-clamp recordings, patch electrodes (Schott 8250 glass, 7-8 MΩ tip resistance for granule cells, 3-4 MΩ tip resistance for Purkinje cells) were filled with a solution containing (in mM): 113 K-Gluconate, 9 HEPES, 4.5 $MgCl_2$, 0.1 EGTA, 14 $Tris_2$-phosphocreatine, 4 $Na_2$-ATP, 0.3 tris-GTP; ~290 mOsm, pH: 7.2-7.25. For voltage-clamp recordings and synaptic activity in Purkinje cells, patch electrodes (Schott 8250 glass, 3-4 MΩ tip resistance) were filled internal solution contained (in mM): 110 $CsMeSO_3$, 40 HEPES, 1 KCl, 4 NaCl, 4 Mg-ATP, 10 Na-phosphocreatine, 0.4 $Na_2$-GTP, 0.1 EGTA; ~290 mOsm, pH: 7.22. All data were corrected for measured junction potentials of 12 and 11 mV in K- and Cs-based internals, respectively.

Electrophysiological data were acquired using Multiclamp 700A or 700B amplifiers (Molecular Devices) via custom routines in IgorPro (Wavemetrics). For measurements of action potential waveform and fiber volleys, data were acquired at 50 kHz and filtered at 20 kHz. For all other measurements, data were acquired at 10-20 kHz and filtered at 3-10 kHz. For current-clamp recordings, pipette capacitance was compensated by 50% of the fast capacitance measured under gigaohm seal conditions in voltage-clamp prior to establishing a whole-cell configuration, and the bridge was balanced. For voltage-clamp recordings, pipette capacitance was compensated completely, and series resistance was compensated 50%.

Parallel fiber volley recordings were made from coronal slices of the medial posterior cerebellum. The recording electrode was placed ~1 mm away from a tungsten bipolar stereotrode (WE3ST30.1A10; MicroProbes for Life Science). The strength of electrical stimulation was adjusted for each recording to maintain the amplitude of the first parallel fiber volley above 1 mV. Parallel fiber volleys were evoked by field electrical stimulation in either trains of 10 at 20 ms inter-stimulus-interval or trains of 20 at 6 ms ISI. This experiment was performed in the presence of 10 μM 6,7-dinitroquinoxaline-2,3-dione (DNQX) and 25 μM picrotoxin.

In experiments measuring paired pulse ratio and synaptic plasticity, EPSCs were evoked via a bipolar glass theta electrode placed ~200 μm orthogonal and ~200 um lateral to the recorded neuron in the molecular layer in the presence of 25 μM picrotoxin at -80 mV.

Whole-cell voltage-clamp recordings were made from the lobule 4/5 of the cerebellum. In short-term synaptic facilitation protocols, excitatory postsynaptic currents (EPSCs) were evoked by field electrical stimulation in a train of 20 at 6 ms ISI.

In high-frequency burst-based plasticity protocols, excitatory postsynaptic currents (EPSCs) were evoked with a theta stimulating electrode placed ~200 μm into the molecular layer and ~200 um lateral to the recorded Purkinje cell. After establishing a stable baseline of 5 minutes (EPSC ISI: 15 seconds) in voltage clamp, EPSPs were evoked by field electrical stimulation in a train of 10 at 6 ms ISI. Trains were repeated every 1 s for 5 minutes. Following induction, EPSC stimulation frequency was reset to 0.05 Hz, and changes in EPSC amplitudes were assessed by comparing data 25 - 30 min following induction to baseline (-5.0 - 0 min).

**Statistics:**

Statistical analyses were performed using IgorPro8 unless otherwise noted (Wavemetrics). No statistical tests were used to predetermine sample size, but our sample sizes are similar to those generally employed in the field. All data are presented in text as mean ± standard error (SEM), unless otherwise noted. The stated P values are the results of the non-parametric Wilcoxon rank sum test to compare values between different mice or recordings, and the non-parametric Wilcoxon signed rank test to compare values from the same recording in different experimental conditions, unless otherwise noted. Mixed-effects modeling was used to compare simple spike firing frequency between baseline and post-induction (units nested within parent animals) using Prism. Data were corrected for multiple comparisons when necessary, using the Holm-Šídák method.

**Supplemental Materials References**
